# Incomplete recruitment of protective T cells facilitates *Trypanosoma cruzi* persistence in the mouse colon

**DOI:** 10.1101/2021.02.26.433032

**Authors:** Alexander I. Ward, Michael D. Lewis, Martin C. Taylor, John M. Kelly

## Abstract

*Trypanosoma cruzi* is the etiological agent of Chagas disease. Following T cell mediated suppression of the acute phase infection, this intracellular eukaryotic pathogen persists in a limited sub-set of tissues at extremely low-levels. The reasons for this tissue-specific chronicity are not understood. Using a dual bioluminescent:fluorescent reporter strain, which allows experimental infections to be imaged at single-cell resolution, we have characterised the ‘hyper-local’ immunological microenvironment of rare parasitized cells in the mouse colon, a key site of persistence. We demonstrate that incomplete recruitment of T cells to infection foci permits the occurrence of repeated cycles of intracellular parasite replication and differentiation to motile trypomastigotes at a frequency sufficient to perpetuate chronic infections. The life-long persistence of parasites in this tissue site continues despite the presence, at a systemic level, of a highly effective T cell response. Overcoming this low-level dynamic host:parasite equilibrium represents a major challenge for vaccine development.

## INTRODUCTION

The insect-transmitted protozoan parasite *Trypanosoma cruzi* is the causative agent of Chagas disease, and infects 5-7 million people in Latin America (1). Despite decades of effort, only limited progress has been made in developing a vaccine, and doubts remain about the feasibility of vaccination as a method of disease control (2,3). In humans, *T. cruzi* infection passes through an acute stage that lasts 2-8 weeks, during which parasitaemia is readily detectable, although symptoms are generally mild and non-specific. With the induction of the adaptive immune response, in which CD8^+^ IFN-γ^+^ T cells play a key role (4,5), there is a significant reduction in the parasite burden. However, sterile clearance is not achieved and parasites persist as a chronic life-long infection. One-third of those infected with *T. cruzi* eventually develop Chagasic pathology, although symptoms can take decades to become apparent. Cardiomyopathy is the most common clinical outcome (6–8), followed by digestive tract megasyndromes, which are reported in about 10% of infected individuals, often in parallel with cardiac disease.

Although the innate immune system is able to detect the parasite (9,10), there is a delay in the subsequent induction of an adaptive response relative to other pathogens (5,11). This, together with a rapid rate of parasite division (12), allows *T. cruzi* to disseminate widely during the acute stage, with most organs and tissues becoming highly infected (13). The CD8^+^ T cell response, which predominantly targets a sub-set of immunodominant epitopes in members of the hypervariable *trans*-sialidase surface antigen family (14,15), is critical for controlling the infection in mice. The parasite burden is reduced by 2-3 orders of magnitude as the disease transitions to a chronic dynamic equilibrium (13). Understanding why the immune system then fails to eliminate the remaining parasites is a central question in Chagas disease research. This information is crucial to underpin rational vaccine design and immunotherapeutic interventions.

Because of the complexity and long-term nature of Chagas disease in humans, mice have been important experimental models for research on interactions between parasite and host. They display a similar infection profile to humans, exhibit chronic cardiac pathology, and are widely used in drug and vaccine development (16). Imaging studies have revealed that the GI tract is a major parasite reservoir during chronic infections and that the degree of containment to this region is determined by both host and parasite genetics (13,17). Parasites are also frequently detectable in the skin, and in some mouse models, skeletal muscle can be an additional site of persistence (4,18). In the colon, the most frequently infected cells are myocytes located in the gut wall. However, the extent of infection is low, and in many cases, this entire organ contains only a few hundred parasites, concentrated in a small number of host cells (18). After transition to the chronic stage, *T. cruzi* also exhibits a reduced proliferation rate, although the cycle of replication, host cell lysis and re-infection appears to continue (12), with little evidence for the type of wide-spread dormancy that characterises other pathogens which establish true latency.

Multiple studies have shown that experimental *T. cruzi* vaccines have protective efficacy and can reduce both parasitaemia and disease severity (19–24). However, unambiguous evidence of complete parasite elimination after challenge, is lacking. In contrast, drug-cured infections can confer sterile long-lasting protection against rechallenge with a homologous parasite strain (3), although this level of protection was only achieved in ~50% of animals. Re-challenge with a heterologous strain did not result in sterile protection, although there was >99% reduction in the peak acute stage parasite burden. All drug-cured animals that displayed re-infection transitioned to the canonical chronic stage equilibrium and organ distribution, without passing through an elevated acute stage parasitaemia. Once established in permissive sites, such as the GI tract, parasites appear to survive the systemic *T. cruzi*-specific IFN-γ^+^ T cell response generated by the primary challenge. In the absence of information on the immunological micro-environment of these persistent parasites, the reasons for this are unclear. Resolving this question will have a major strategic impact on the development of an effective vaccine.

Progress in this area has been limited by technical difficulties in locating and analysing the rare infection foci in permissive tissue sites, such as the colon. Here, we describe the application of a *T. cruzi* bioluminescent:fluorescent dual reporter strain and enhanced imaging procedures that have allowed us to show that incomplete ‘hyperlocal’ homing of T cells to foci of intracellular infection is associated with the ability of the parasite to persist in the colon.

## Results

### Cellular immunity suppresses the colonic parasite load during chronic *T. cruzi* infection

Myocytes in the colonic gut wall are an important site of *T. cruzi* persistence in murine models of chronic Chagas disease. However, infected host cells are extremely rare and unevenly distributed (18). To assess the role of the cellular immune response in controlling infection in this tissue compartment, we infected C3H/HeN mice with *T. cruzi* CL Luc::mNeon, a parasite line that constitutively expresses a bioluminescent:fluorescent fusion protein (25). This reporter strain can be used in combination with *ex vivo* imaging and confocal microscopy of colonic wall whole mounts to detect persistent infection foci at single cell resolution (Materials and Methods). When infections had reached the chronic stage (>100 days post-infection), one cohort of mice was immunosuppressed with cyclophosphamide, an alkylating agent that is generally suppressive of the lymphocyte population (26), and which has been widely used to drive the reactivation of low-level *T. cruzi* infections (27,28). Treatment led to a major reduction in peripheral blood mononuclear cells (PBMCs) within 5-10 days (Figure 1a, b). In parallel, other groups of mice were subjected to antibody-mediated depletion of the circulating CD4^+^ or CD8^+^ T cell populations. This was achieved, with high specificity, in a similar time-scale (Figure 1c, Figure 1 - figure supplement 1). Circulating anti-*T. cruzi* serum antibody levels were not significantly altered by cyclophosphamide treatment, or by depletion of the CD4^+^ or CD8^+^ T cell subtypes (Figure 1d).

**Figure 1.**
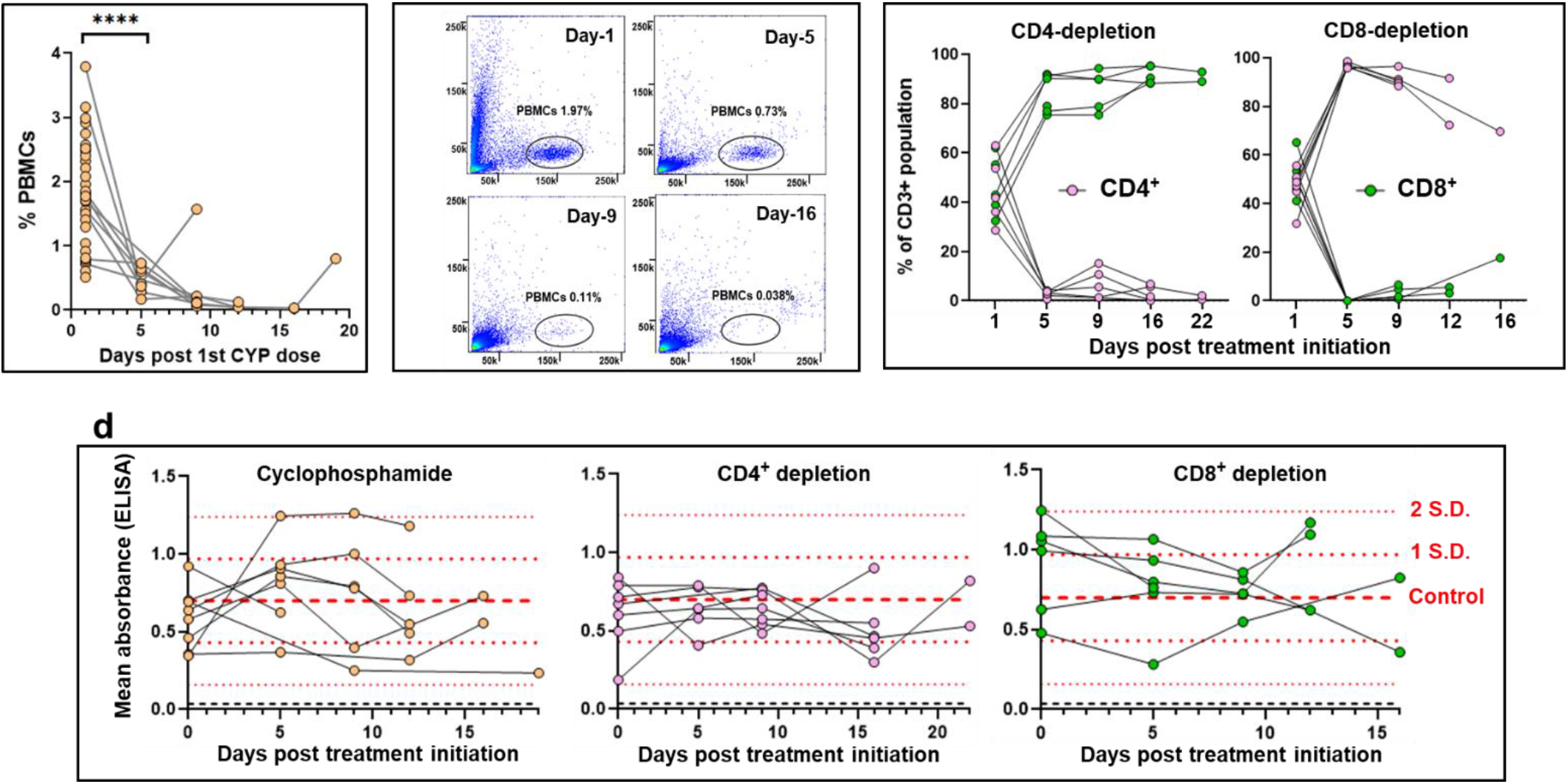
Suppression of cellular immunity in mice chronically infected with *T. cruzi.* **a.** C3H/HeN mice chronically infected (>100 days) with *T. cruzi* CL Luc::mNeon (n=6) were immunosuppressed by i.p. inoculation with cyclophosphamide (200 mg/kg) at 4-day intervals, up to a maximum of 3 injections (Materials and Methods). The % events recorded as peripheral blood mononuclear cells (PBMCs) at different time points after the initiation of treatment for individual mice are shown (Materials and Methods). Also included in the day 1 values are additional data points (n=24) from immunocompetent chronically infected mice. **b.** Flow cytometry plots showing the loss of detectable events in the PBMC gate (black oval) over the course of cyclophosphamide treatment (see also Figure 1 - figure supplement 1) PBMCs were identified based on the spectral forward (FFC, Y-axis) and side (SSC, X-axis) scatter. **c.** Effective depletion of T cell subsets by treatment of mice with specific anti-CD4 or anti-CD8 antibodies (Materials and Methods). The graphs show the CD4^+^ and CD8^+^ flow cytometry events of individual mice as a % of the total CD3^+^ population over the treatment periods. **d.** ELISA mean absorbance readings (using anti-mouse IgG secondary antibody) for serum from chronically infected mice that had been treated with cyclophosphamide, or treated with anti-CD4 or anti-CD8 antibodies. Microtitre plates containing *T. cruzi* lysates were prepared as described (Materials and Methods). Dashed red lines identify the mean, +1 × S.D. and +2 × S.D. values, determined from immunocompetent chronic stage controls (n=28). One of the anti-CD8 antibody treated mice died between day 5 and 9.

Examination of mouse colons by *ex vivo* bioluminescence imaging >12 days after the initiation of treatment, revealed that cyclophosphamide-induced immunosuppression had resulted in dissemination and increased intensity of the infection (Figure 2a). Further analysis of peeled external gut walls by confocal microscopy (Materials and Methods), which allows the full length of the longitudinal and circular smooth muscle layers of the colon to be assessed at a 3-dimensional level (18), confirmed that there had been a significant increase in the number of infected cells (Figure 2b, c, d). Therefore, cyclophosphamide treatment perturbs the ability of the immune system to control the proliferation of persistent parasites within the colon. However, specific depletion of either the CD4^+^ or the CD8^+^ T cell repertoires by themselves, did not have a significant effect (Figure 2d). Furthermore, in the absence of PBMCs, it is implicit from the resulting parasite dissemination that the circulating serum antibodies are unable to suppress the infection at this site during the chronic stage (Figure 1d).

**Figure 2.**
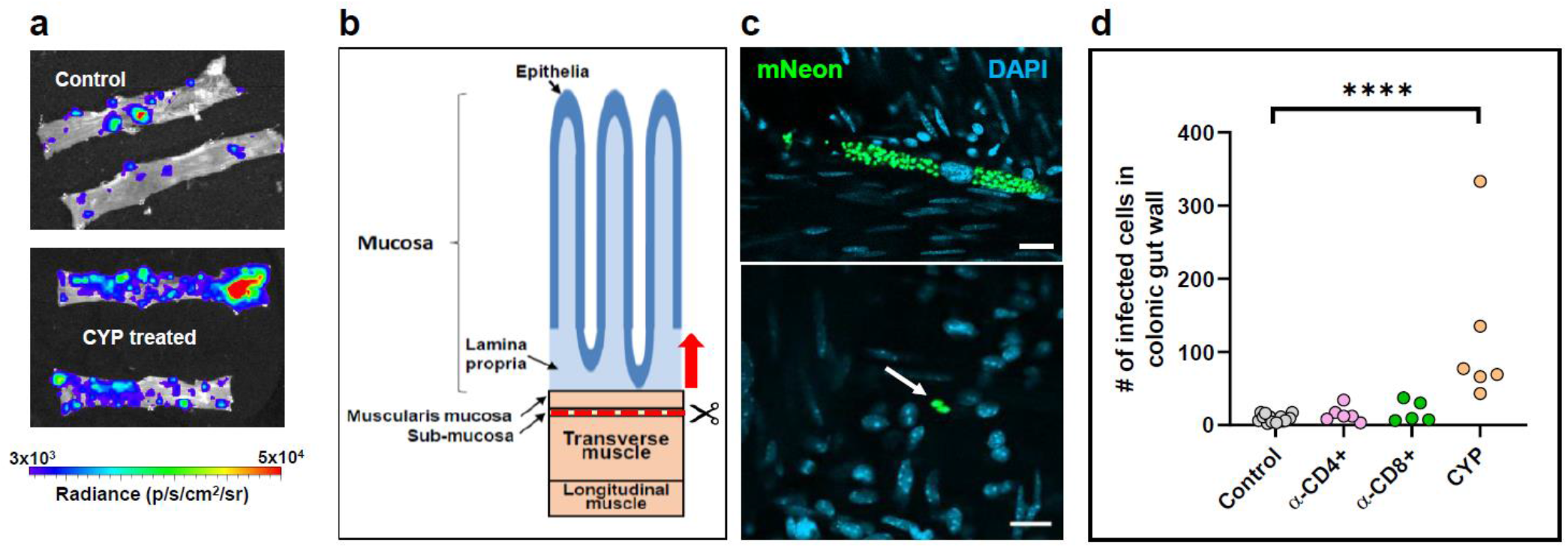
Control of persistent parasites in the colon of chronically infected mice is lost on suppression of cellular immunity. **a.** Colon sections from C3H/HeN mice chronically infected with *T. cruzi* CL-Luc::mNeon were pinned luminal side up and examined by e*x vivo* bioluminescence imaging. Radiance (p/s/cm^2^/sr) is on a linear-scale pseudo-colour heat map. Upper inset, colonic sections from non-treated infected mice; lower inset, section from mice immunosuppressed by cyclophosphamide treatment (Materials and Methods). **b.** Schematic highlighting the distinct layers of the GI tract. The dashed red line and arrow indicate the position above which tissue can be peeled off to leave the external colonic wall layers (18). **c.** External gut wall whole mounts were examined in their entirety at a 3-dimensional level by confocal microscopy. Examples of parasite infected cells and their locations, detected by green fluorescence (mNeon). DAPI staining (blue) identifies host cell nuclei. Scale bars=20μm. **d.** The total number of parasitized cells counted in each whole mounted colonic gut wall for the control and the immune-depleted groups. Each dot represents a single mouse, with the colons examined >12 days post treatment initiation. **** = p≤0.0001. Differences between control values and those obtained from mice that had been treated with anti-CD4 and anti-CD8 antibodies were nonsignificant.

### Parasites persisting in the colon can induce effective hyper-local T cell recruitment

At any one time, the majority of the parasite population that persists in the colon is found in a small number of ‘mega-nests’, infected cells that typically contain several hundred replicating amastigotes, or occasionally, differentiated nondividing trypomastigotes (12). The remainder of the population is more widely distributed, with considerably lower numbers of parasites per infected cell. To better understand the process of long-term parasite survival, we investigated the cellular microenvironment of persistent infection foci. When infections had advanced to the chronic stage, peeled colonic wall whole mounts were examined by confocal microscopy (Materials and Methods), and compared to those of naïve age-matched mice. In the tissue from non-infected mice, using DAPI staining to highlight nuclei, an average of 55 host cells were identified in 200 μm diameter circles positioned around randomly selected nuclei within the whole mounted gut wall (Figure 3a). Most cells had elongated nuclei typical of smooth muscle myocytes. In the infected group, parasitized cells were identified by green fluorescence (Materials and Methods). Scanning revealed that total cellularity in the immediate locality of infection foci was similar in most cases to that in non-infected colon tissue; 95% were within 3 × S.D. of the background mean, compared with 98% around randomly selected cells from naïve control regions (Figure 3a, b). However, on occasions there was evidence of highly localised cellular infiltration, with 3.4% of infection foci surrounded by a local cellularity that was >4 × S.D. above the background mean. Within these intense infiltrates, host cells with more rounded nuclei predominated. In contrast to the majority of parasitized cells that had not triggered a detectable hyper-local immune response (Figure 3c), amastigotes in these inflammatory infiltrates frequently displayed an irregular morphology that suggested immune-mediated damage, as judged by the diffuse pattern of green fluorescence (compare Figure 3c, d, e).

**Figure 3.**
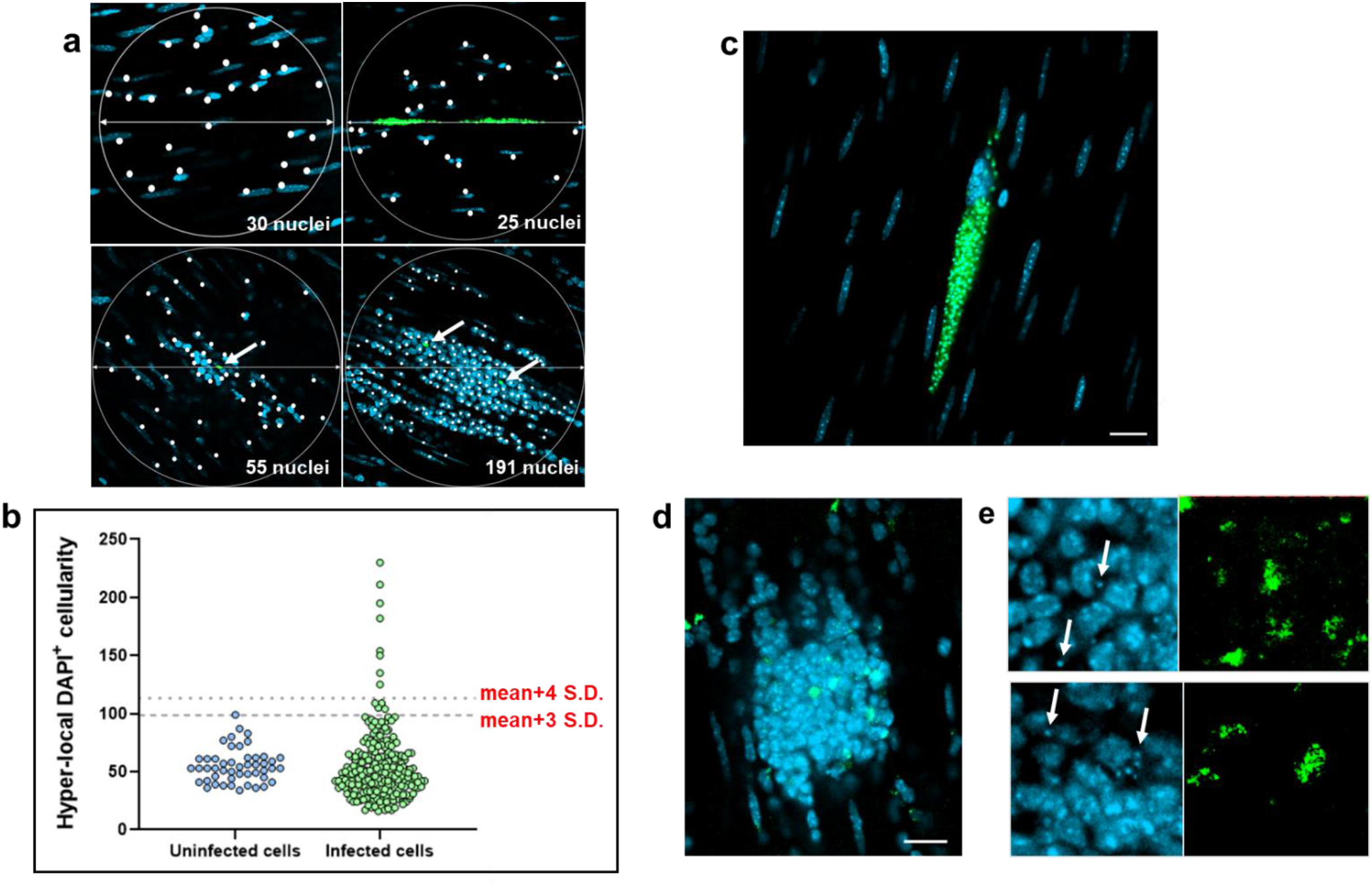
Defining the hyper-local cellularity of *T. cruzi* infected host cells in the colonic gut wall. **a.** Images of whole mounted colonic gut wall from C3H/HeN mice chronically infected with *T. cruzi* CL-Luc::mNeon (Materials and Methods). When infection foci were identified, 200 μM diameter circles were drawn centred on each parasite cluster or ‘nest’. Circles were placed by centering on randomly selected cells in the case of non-infected age-matched controls (top left panel). DAPI-stained nuclei (blue) that fell within this disc (highlighted by white dots) were counted as a measure of cellularity. Intracellular parasites can be identified by green fluorescence. These are indicated by white arrows in the lower images. **b.** Background cellularity around randomly selected cells (n=48) on whole mounted colonic gut walls from naïve age-matched C3H/HeN mice was established as above. With tissue from chronically infected mice, hyper-local cellularity was calculated using circles centred on parasite foci (green) (n=247). Individual values are indicated by blue (non-infected) and green (infected) dots. The dashed lines indicate 3 × S.D. and 4 × S.D. above the background mean. **c.** An infected myocyte where the local cellularity is equivalent to the background level and the intracellular amastigotes (green) are structurally intact. Scale bar=20 μm. **d.** Image of an intense cellular infiltrate (nuclei, blue) in which the *T. cruzi* parasites (green) display an irregular and diffuse morphology. **e.** Zoomed-in views of two regions of the same cellular infiltrate. Many discrete disc-like kDNA structures (the parasite mitochondrial genome network) are detectable by DAPI-staining throughout this inflammatory focus (examples indicated by white arrows). They often co-localise with diffuse green fluorescence of parasite origin (right-hand images).

We investigated the nature of these cellular infiltrates, by staining colonic gut wall whole mounts from chronically infected mice with specific immune cell markers (Materials and Methods). This revealed, as expected, that leukocytes (identified by anti-CD45 antibodies) constituted close to 100% of the infiltrate population (Figure 4a). A major proportion of the recruited cells were also positive when stained with anti-CD3 antibodies, specific markers for the T-cell receptor complex (Figure 4b, c), with both CD4^+^ and CD8^+^ T cells represented within this population (Figure 4d). To assess the local density of stained immune cells, we examined 200 μm diameter circular tissue sections centred on each infection focus using Z-stack confocal microscopy. A series of imaged sections starting 5 μm above and 5 μm below the centre of the parasite nest (a total volume of 314 μm^3^) were generated, and the number of stained cells in the infection microenvironment determined in 3-dimensions (Figure 4 - supplement figure 1). In sections of colonic smooth muscle from non-infected mice, leukocytes were dispersed and rare, with an average of ~1 CD45^+^ cell per 314 μm^3^, although they were more numerous in the sub-mucosal tissue (Figure 4 - supplement figure 2). Using a cut-off value of 3 × S.D. above the respective background level, 40 - 45% of infection foci displayed evidence of leukocyte infiltration (Figure 4e). Therefore, despite being a site of parasite persistence, dynamic hyper-local homing of T cells to parasitized cells in the murine colon is a characteristic of chronic stage infection, although at any one point in time, not all parasite nests will have triggered this type of recruitment response. Given the ‘snapshot’ nature of imaging, our data therefore suggest that in the majority of cases, the most likely outcome of colonic cell invasion will be infiltration of leukocytes, and the presumptive destruction of the parasites (Figure 3d, e).

**Figure 4.**
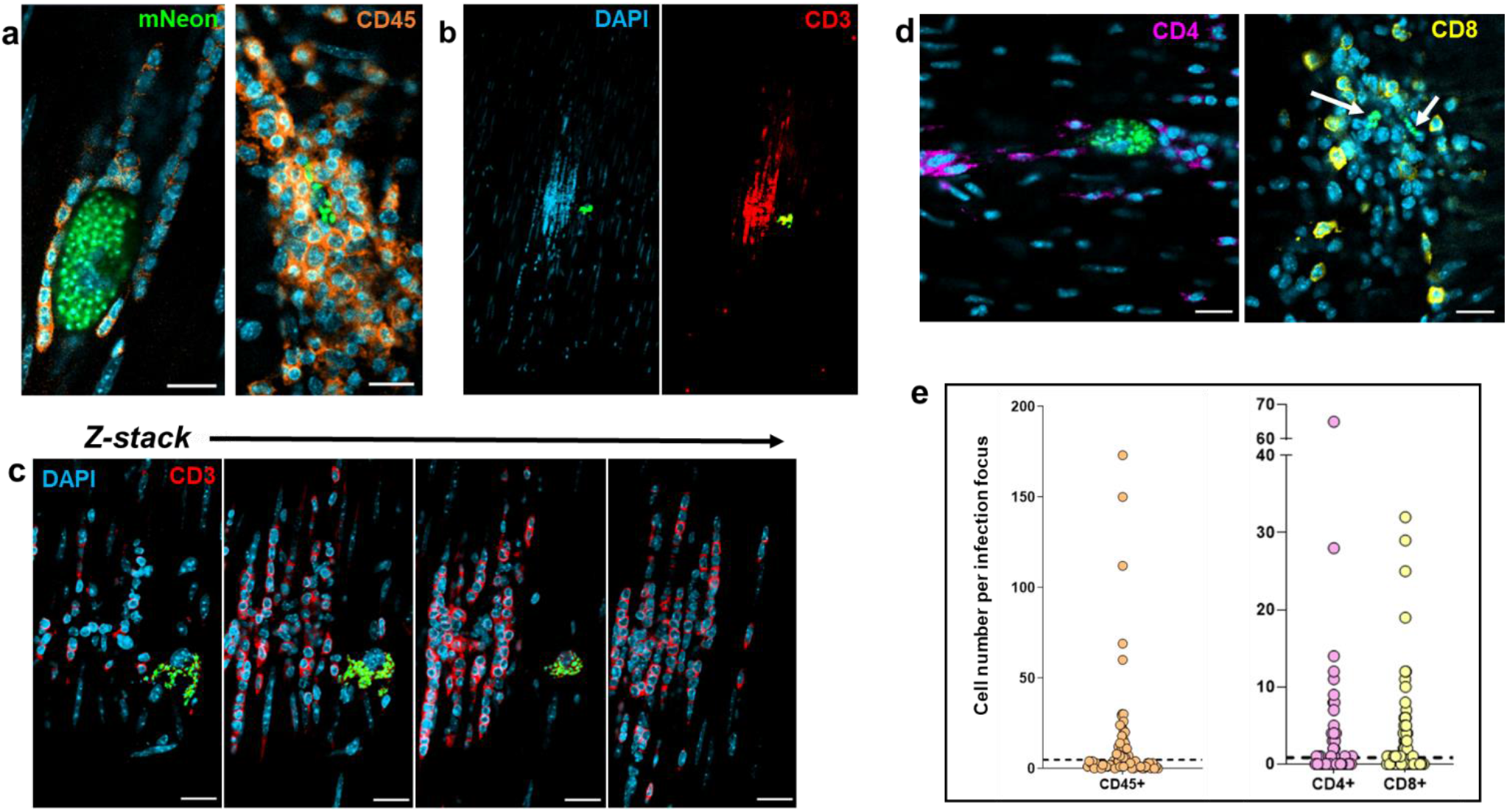
T cells are major constituents of the leukocyte population recruited to chronic stage infection foci. **a.** Confocal images of colonic gut wall sections from chronically infected mice (Materials and Methods). Rare infection foci were identified by mNeonGreen fluorescence (parasites) after exhaustive searching of whole mounted gut walls. Staining with anti-CD45 (orange) reveals that hematopoietic cells constitute the vast majority of the infiltrate population. Host cell nuclei were identified by DAPI staining (blue). **b.** Anti-CD3 staining of cellular infiltrates shows that T-cells constitute a majority of the population. Blue, host cell nuclei; red, CD3 staining; green, parasite fluorescence. **c.** Serial Z-stack imaging (Materials and Methods) through the same cellular infiltrate as in b, showing selected sections through the infiltrate. **d.** Histological sections containing cellular infiltrates and associated infection foci (parasites, green; indicated by white arrows in right-hand image) stained with either anti-CD4 (purple) or anti-CD8 (yellow) antibodies. Scale bars=20 μm. **e.** Whole mounts containing infection foci were stained with anti-CD45, anti-CD4, or anti-CD8 antibodies and the number of positive host cells in the immediate vicinity (314 μm^3^ volume) was determined by serial Z-stack confocal imaging (see also Figure 2). Each dot corresponds to a single infection focus. The horizontal dashed line is 3 × above the S.D. of the mean background level in non-infected tissue. In the case of anti-CD45 staining, none of the 50 tissue sections examined from non-infected mice contained CD45+ve positive cell numbers above this value. 41%, 45% and 42% of infection foci identified by CD45, CD4 and CD8 staining, respectively, were above this cut-off.

### Incomplete homing of protective T cells allows a subset of intracellular colonic infections to complete their replication cycle

Evidence indicates that *T. cruzi* rarely occupies individual colonic myocytes for extended periods (>2 weeks) (12), implying that parasites are either efficiently eliminated by the immune response, or that they complete a cycle of replication, trypomastigogenesis and host cell lysis within this period. In addition, there is considerable variation in the level of infection within individual colonic cells, with parasite numbers that can range from 1 to >1000 (12). We therefore investigated whether the immune response induced against infected cells increased in line with the intra-cellular parasite burden. When the levels of infiltrating leukocytes in the local environment of infected cells were compared with the number of intracellular *T. cruzi* parasites, we found no apparent correlation (Figure 5a, b, c). This was the case irrespective of whether anti-CD45, anti-CD4 or anti-CD8 antibodies were used to assess the nature of the cellular infiltrate. It is implicit therefore, that the elapsed duration of an individual intracellular infection, as inferred from the extent of parasite proliferation, is not a determinant of the likelihood of detection and targeting by the host immune system.

**Figure 5.**
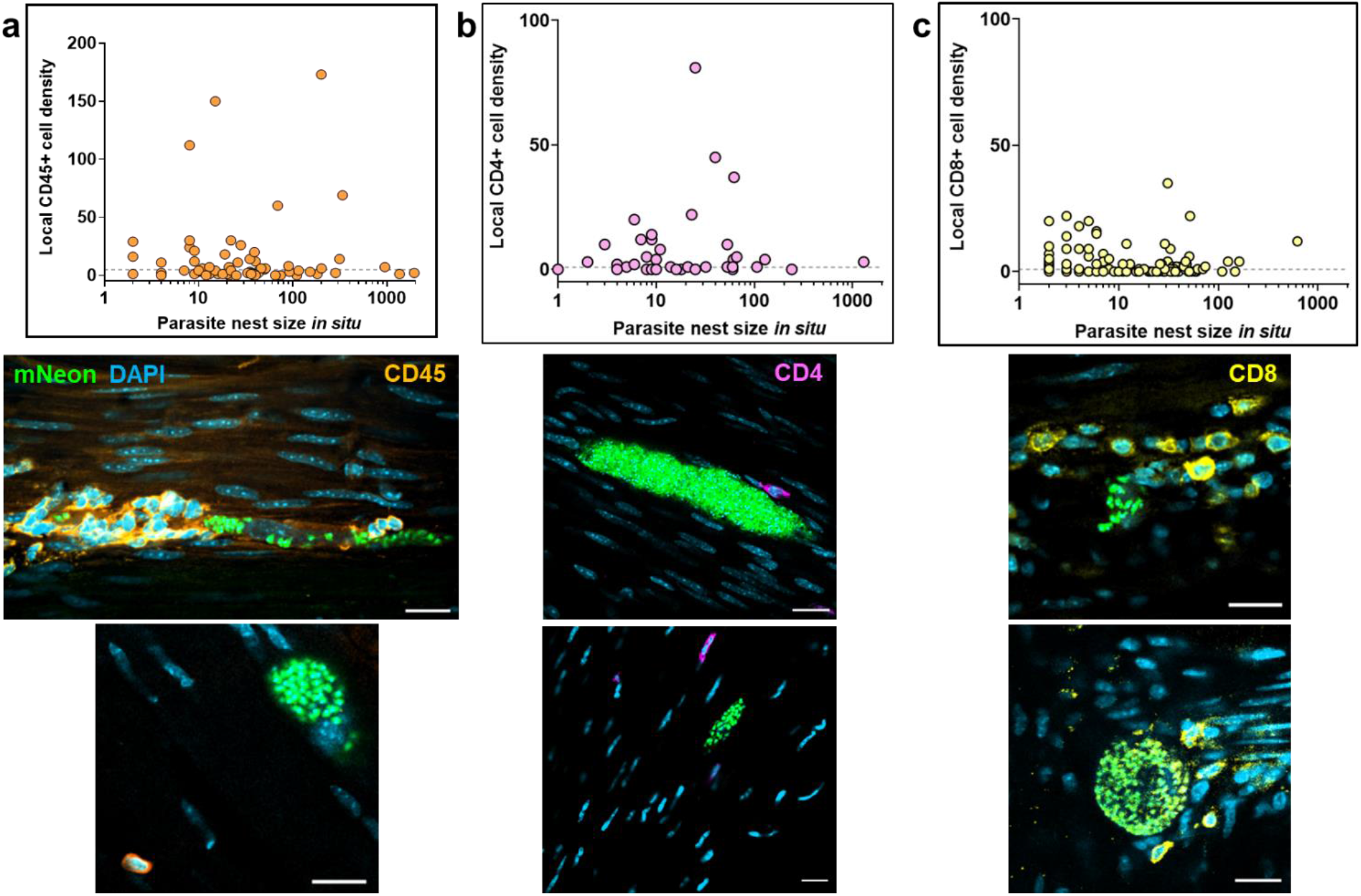
Lack of correlation between intracellular parasite load and hyper-local T cell infiltration during chronic infections. **a.** Comparison of the parasite numbers in infected colonic gut wall cells with the local leukocyte cell density. Infection foci were identified in whole mounts of colonic tissue, which were then stained with anti-CD45 antibody (Materials and Methods). The parasite and cell numbers in a tissue volume of 314 μm^3^ were determined using serial Z-stack imaging, with leukocytes identified by orange staining and parasites by green fluorescence. The horizontal dashed line is 3 × above the S.D. of the mean background level in non-infected tissue. Each dot identifies a single infection focus, with tissue samples derived from 6 mice (71 infection foci). The confocal images show representative infection foci used to generate the data, and illustrate the varying extents of leukocyte infiltration. **b.** Similar analysis of infection foci using anti-CD4 staining (purple). Tissue samples were derived from 3 mice (54 infection foci). **c.** Analysis of infection foci using anti-CD8 staining (yellow). Tissue derived from 4 mice (116 infection foci).

Of 237 infected colonic cells detected in 13 animals, only 4 (~1.7%) contained parasites that had clearly differentiated into flagellated trypomastigotes, the life-cycle stage that disseminates the infection by re-invasion of other host cells, or via transmission to the blood-sucking triatomine insect vector. Of these, three contained large numbers of parasites (>1,000), while the fourth contained 128. In each case, the leukocyte densities in the local microenvironment were within a range similar to host cells where the infection was less advanced, as judged by the number of intracellular parasites and the lack of differentiation into trypomastigotes. In the example shown (Figure 6a, b), Z-stack imaging was used to serially section a mega-nest containing >1,000 parasites, and shows mature trypomastigotes in the act of egress, despite the recruitment of a small number of CD45^+^ leukocytes, including CD8^+^ T cells (Figure 6c, d). Although the precise signals that trigger differentiation to the trypomastigote stage are unknown, it can be inferred from our data that the differentiation process itself does not act to promote rapid infiltration of leukocytes to the site of infection, at least in the colon. Therefore, for a small proportion of infected cells, the host immune system is either not triggered locally by an infection, is too slow to respond, or is in some way blocked. As a result, in the colon, the entire cycle of parasite proliferation, differentiation and egress can occur in the absence of intervention by a cellular immune response, leading to the invasion of new host cells and prolongation of the chronic infection.

**Figure 6.**
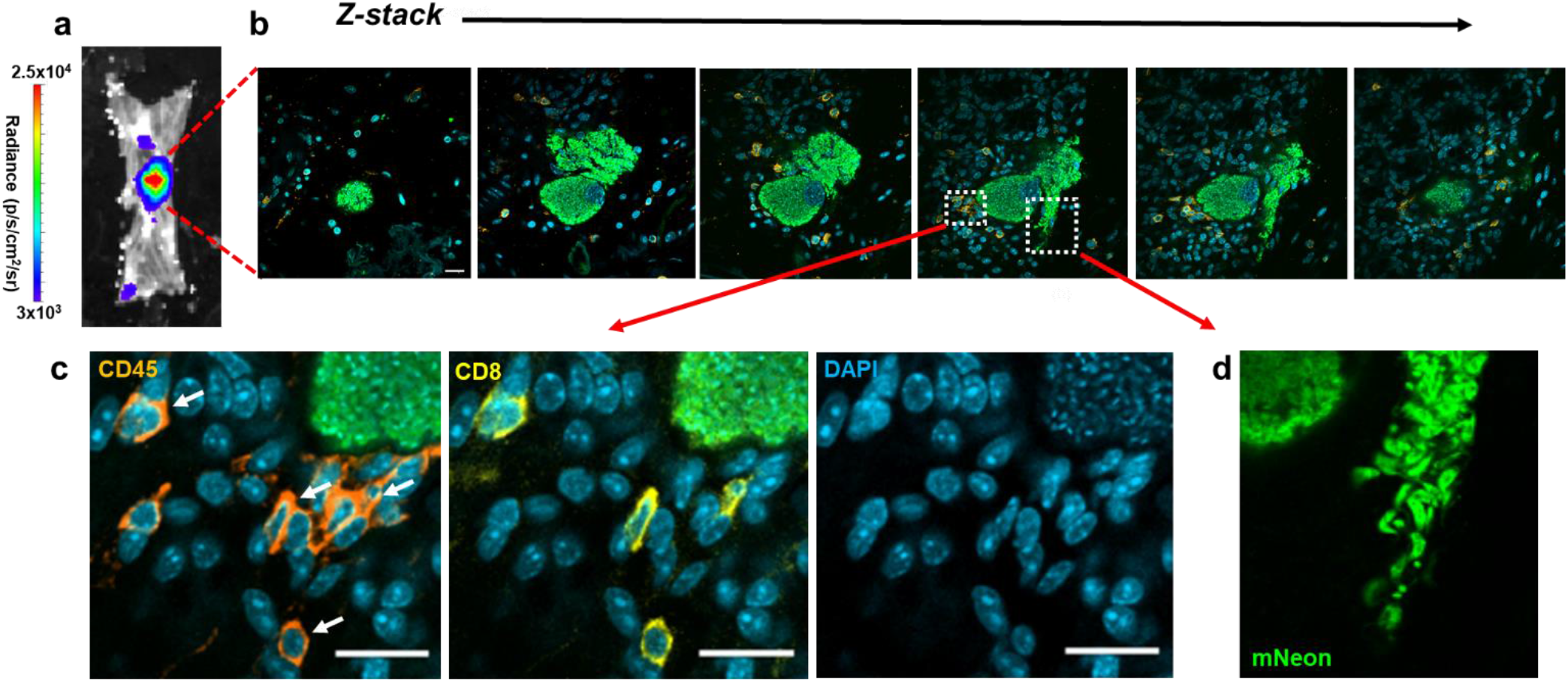
Incomplete recruitment of leukocytes allows progression of *T. cruzi* through the full intracellular infection cycle. **a.** An intense bioluminescent focus in a chronic stage distal colon viewed by *ex vivo* imaging (Materials and Methods). Radiance (p/s/cm^2^/sr) is on a linear-scale pseudocolour heatmap. **b.** Confocal imaging of the corresponding parasite mega-nest showing representative serial Z-stack images taken along the depth of the infected cell. The Z-axis position relative to the centre of the nest is indicated above each of the images. Parasite numbers (>1000) were established from green fluorescence and the characteristic DAPI staining of the parasite kinetoplast DNA (the mitochondrial genome) (18) (blue). Infiltrating leukocytes (orange) were identified by staining with anti-CD45 antibodies (Materials and Methods). Scale bar=20 μm. **c.** Enlarged images of a small cluster of infiltrating CD45^+^ (orange) and CD8^+^ (yellow) cells in close vicinity to the nest. White arrows indicate leukocytes corresponding to CD8^+^ T cells. **d.** Egress of differentiated trypomastigotes into the extracellular environment. Data from the infected cell captured in these images was not included in Figure 5 since the parasite burden was too great to determine with precision.

## Discussion

Despite the generation of a vigorous and specific CD8^+^ T cell response (4,14,29,30), *T. cruzi* infections in mice are rarely cleared to sterility, even in vaccinated animals. Instead, the parasite persists in a small number of reservoir tissue sites, typically for the life-time of the host (10). Intermittent dissemination from these locations to less permissive organs, such as the heart, may promote repeated episodes of infection, resulting in localised inflammatory responses that contribute to disease pathology in a cumulative manner (31). Understanding why the immune system efficiently suppresses, but fails to eliminate *T. cruzi* infections, is one of the key challenges in Chagas disease research. Here, using techniques that allow the immunological microenvironment of infection sites to be assessed at single cell resolution, we demonstrate that both CD4^+^ and CD8^+^ T cells are frequently recruited to chronic infection foci in the colon, and that parasites in this site of persistence are subject to immune-mediated destruction. However, for a small sub-set of infected cells, recruitment is either absent, or too slow, to prevent completion of the intracellular cycle of parasite proliferation and differentiation to the trypomastigote stage (Figure 6). Thus, chronic *T. cruzi* infections in the colon are not characterised by a generalised tissue-specific latency, but by a dynamic equilibrium between host and pathogen.

T cell recruitment during *T. cruzi* infection is driven by secretion of chemokines from infected cells. For example, the CXCR3 ligands CXCL9 and CXCL10 have been implicated in cardiac infiltration (32). IFN-γ and TNF-α expression by antigen specific CD8^+^ T cells (4), and subsequent iNOS expression (33–35), potentially from recruited innate monocytes or from somatic cells of the infected tissue, then increases the local concentration of reactive nitrogen species. In Chagas disease, the resulting inflammatory environment tightly controls the number of infected cells, but can also act as the key driver of chronic immunopathology (7,14,36,37). An important observation from our study is that the likelihood of T cell recruitment is not linked with the maturity of individual *T. cruzi* infections, as judged by the intracellular parasite load (Figure 5). In addition, the process of differentiation to the flagellated trypomastigote form, which occurs in highly parasitized cells in an asynchronous manner (38), does not appear to be a key trigger that enhances infiltration of leukocytes, including CD8^+^ cells, to the site of infection (Figure 6).

The reasons why protective T cells are not recruited to a small sub-set of infection foci are unclear. Hypothesised mechanisms to account for *T. cruzi* immune evasion include a general absence of pathogen associated molecular patterns (PAMPs) (39), cytokine-mediated inhibition of effector responses (10), insufficiently strong chemoattractant signalling in low parasite load settings (36), the extensive antigenic diversity expressed by the large families of *trans*-sialidase and mucin genes (14,40,41), and stress-induced cell-cycle arrest and dormancy (42). However, none of these obviously correspond with our observation that there is an apparent lack of association between the extent or longevity of an individual cellular infection and the magnitude of hyper-local leukocyte recruitment (Figure 5). Some highly infected meganests appear to be invisible to the immune system, whereas other low-level infections trigger massive cellular infiltration. One explanation could be that a slow-down in the intracellular amastigote replication rate during chronic stage infections (12) contributes to reduced immune detection. In circumstances where the infected cell is in an area of the colon that is otherwise parasite-free, this may be sufficient to permit completion of the initial replication cycle. However, after trypomastigote egress and host cell lysis, the resulting tissue disruption and production of damage associated molecular patterns (DAMPs) could act to enhance T cell recruitment into the area, leading to the destruction of parasites that have re-invaded host cells in the vicinity of the initial infection. In contrast, trypomastigotes which migrate away from this locality may be able to establish a productive infection in the absence of rapid immune detection. Despite a diverse and complex antigenic repertoire, induction of the T cell response in draining lymph nodes is known to be highly focussed (14), and once T cell recruitment has been triggered, parasite destruction can be initiated (Figure 3d, e). Widespread parasite dormancy was not evident in colonic tissue (12), and does not appear to be necessary for immune evasion.

Success or failure of the immune system in eliminating these rare chronic infection foci may be a largely stochastic process resulting from the dynamic interplay between the host and pathogen at a single cell or tissue micro-domain level. If parasites were able to universally suppress innate detection pathways, with concomitant reduction in localised chemokine output, this would have a negative impact on host survival, and thus long-term *T. cruzi* transmission. Conversely, if infections were always detected by the immune system before completion of the replication cycle, the parasite would risk host-wide elimination. The ability of *T. cruzi* to persist in some organs/tissues, may therefore be dependent on the propensity, or otherwise, of these tissues to amplify the chemokine signals triggered by low-level infection, with a possible role for closely adjacent re-infections in the amplification process. In mice, there are strain-specific differences in the extent of such tissue-restriction during chronic infections. This could have parallels in humans, and account for the heterogeneous profile of disease progression.

*T. cruzi* infection induces a high titre polyclonal B cell/antibody response during the acute stage of infection, which although delayed and initially unfocussed (43), does contribute to parasite control and can protect against virulent infections. In the chronic stage, a role for the humoral response in suppressing the dissemination of persistent parasites is unresolved (10), and a key role for B cells has not been identified. Here, we show that in the absence of PBMCs, circulating antibodies, which in the short-term are not profoundly affected by cyclophosphamide treatment (44) (Figure 1d), are unable to maintain tissue-specific repression of the parasite burden (Figure 2d). If the humoral response does have a significant protective role during the chronic stage, for example, involving opsonisation of the parasite through FcR-antibody binding, then this function could be lost on depletion of key cellular effectors. In addition, our results do not exclude the possibility that parasite-specific antibodies could act to limit infections at a systemic level, over a longer duration, perhaps by controlling trypomastigote numbers.

The central role of CD8^+^ T cells in suppressing *T. cruzi* infections is well established, and in various parasite:mouse strain combinations, depletion of circulating CD8^+^ T cells leads to partial recrudescence, at least in skeletal muscle and adipose tissue (4,5,15,30). When we examined the effect of CD8^+^ T cell depletion at a cellular level in colonic tissue, we found no significant increase in the number of infected cells, in contrast to the major rebound observed with cyclophosphamide-mediated reduction of the entire PMBC population (Figure 1,2). A non-redundant function for CD4^+^ T cells is less well established in murine models of Chagas disease (45–47), although in humans with untreated HIV co-infections, parasites become easily detectable in the bloodstream (48). Since depletion of either CD4^+^ or CD8^+^ T cells by themselves did not promote the level of relapse observed with cyclophosphamide treatment over the time period analysed (Figure 2), our results therefore suggest that both lymphocyte sub-types are able to suppress chronic stage infections in the colon, together with innate leukocytes that may mediate and enhance T cell effector functions. Similar observations have been made using PCR-based detection to examine skeletal muscle of C57BL/6 mice infected with the *T. cruzi* Colombiana strain (37).

Our findings have important implications for anti-*T. cruzi* vaccine development. Vaccines protect by presenting non-tolerised antigens in the correct immunological context, to expand small numbers of antigen-specific naïve T and B cells, which then generate a sub-population of memory cells. The expanded memory populations then allow more rapid deployment of adaptive effectors on future contact with the pathogen. However, *T. cruzi* is able to persist indefinitely in hosts that already have expansive systemic populations of effective T cells. Unless vaccines can prevent parasites from accessing sites of persistence after the initial infection, or they are able to enhance successful homing of adaptive effector cells, it will be difficult to achieve sterilising immunity. Drug-cured infections can confer complete protection against re-challenge with a homologous strain, but with heterologous strains, despite the prevention of an acute stage peak, the infection proceeds directly to a status that is analogous to the chronic stage in terms of parasite burden and tissue distribution (3). Therefore, it is likely that successful anti-*T. cruzi* vaccines will require an ability to eliminate parasites at the initial site of infection during the first intracellular replication cycle. This will be a considerable challenge.

## Materials and Methods

### Mice and parasites

All experiments were performed using female C3H/HeN mice, purchased from Charles River (UK). They were maintained in individually ventilated cages, under specific pathogen-free conditions, with a 12-hour light/dark cycle, and provided with food and water *ad libitum.* Research was carried out under UK Home Office project licenses PPL 70/8207 and P9AEE04E4, with approval of the LSHTM Animal Welfare and Ethical Review Board, and in accordance with the UK Animals (Scientific Procedures) Act 1986 (ASPA). The *T. cruzi* line CL Luc::mNeon, a derivative of the CL Brener strain (discrete typing unit TcVI), was used in all experiments. It had been genetically modified to express a bioluminescent:fluorescent fusion protein containing red-shifted luciferase and mNeonGreen fluorescent domains (25,49). For infections, C3H/HeN mice, aged 6-8 weeks, were inoculated i.p. with 1×10^3^ bloodstream trypomastigotes obtained from immunodeficient CB17-SCID mice, as described previously (28). Mice were then monitored by *in vivo* bioluminescence imaging (17) which indicated that they had transitioned to the chronic stage by 50-60 days post-infection. Experiments were performed when mice had been infected for at least 100 days.

### Suppression of the murine immune response

General immunosuppression was achieved by injecting mice i.p. with cyclophosphamide (200 mg/kg) at 4-day intervals, up to a maximum of 3 injections, in accordance with animal welfare (17,28). Circulating CD8^+^ T cells were depleted by i.p. injection of 400 μg of the YTS 169.4 monoclonal anti-CD8α (2BScientific), diluted in PBS, at 4-day intervals, up to a maximum of 4 times. The same regimen was applied for depletion of CD4^+^ T cells, using the GK1.5 monoclonal antibody (2BScientific).

### Tissue processing and imaging

When mice were sacrificed, organs and tissues were removed and examined by *ex vivo* bioluminescence imaging using the IVIS Spectrum system (Caliper Life Science) and the LivingImage 4.7.2 software (50). Colonic muscularis walls were isolated by peeling away the mucosa, whole mounted as described previously (18), and then exhaustively searched for parasites (green fluorescence) with a Zeiss LSM880 confocal microscope. Small tissue sections (~5 mm^2^) around parasite nests were excised from the whole mount by scalpel, washed twice in PBS and incubated for 2 days in 1:300 primary antibody diluted in PBS / 5% fetal calf serum / 1% Triton-X100 at 4°C. Following 2 further washes in PBS, secondary antibody diluted in 1:500 in the same blocking/permeabilising solution was added to the tissue sections, and incubated for 3 hours at room temperature. Sections were then mounted in Vectashield, containing the DNA stain DAPI, and imaged by confocal microscopy. Colonic muscularis walls from naïve aged-matched mice were similarly prepared as controls, with and without the primary antibody.

For accurate determination of intracellular parasite and surrounding host cell numbers, tissue samples were imaged in 3-dimensions (Z-stacking), with the appropriate scan zoom setting (18). The Image Browser overlay function was used to add scale bars, and images were exported as .TIF files to generate figures. Primary antibodies used were as follows: anti-luciferase (G7451, Promega), CD45 (Tonbo Biosciences, 30- F11), CD3 (Abcam, ab11089), CD4 (Abcam, ab25475), CD8 (Abcam, ab25478). The secondary antibodies were Invitrogen A-11055, Invitrogen A-21434, Invitrogen A- 11007.

### Flow cytometry

At each time-point, mice were placed in a “hot box” and left at 38°C for 10 minutes. They were then placed in a restrainer and the lateral tail vein punctured using a 0.5M EDTA (pH 7.4) soaked 21G needle. A single drop of blood was transferred to a 2 ml tube and 10μl 0.5M EDTA added to prevent clotting. Each sample was then mixed with 400 μl ice-cold PBS and placed onto 300 μl Histopaque 1083 (Sigma-Aldrich), and spun at 400 *g* for 30 minutes in a microcentrifuge. The monocytic layer was aspirated using a pipette, mixed with 1 ml ice-cold PBS, pelleted and resuspended in 200 μl flow cytometry buffer (PBS, 5% fetal bovine serum, 0.05% sodium azide), and 1 μl of the cocktail of conjugated antibodies added (1:200 dilution in each case). After 1 hour incubation in the dark, cells were pelleted and resuspended in 2% paraformaldehyde in PBS, followed by a further 45 minutes incubation in the dark. The stained fixed cells were then pelleted, re-suspended in filtered flow cytometry buffer and transferred to standard flow cytometry tubes. Samples were analysed using a BD Bioscience LSRII flow cytometer, with plots created and analysed in FlowJo V.10.6.1. The following antibodies were used: CD45 (ThermoFisher, 30-F11, Super Bright 600), CD3 (ThermoFisher, 17A2, FITC), CD4 (ThermoFisher, RM4-5, eFluor 450), and CD8 (ThermoFisher, SK1, Alexa Fluor 780).

### *α-T. cruzi* antibody ELISA

96-well plates were coated with sonicated *T. cruzi* CL Luc::mNeon trypomastigote lysate; 100 μl (0.5 μg) per well diluted in 15 mM Na2CO3, 34.8 mM NaHCO3. The plates were incubated at 4°C overnight to allow antigen binding, washed 3x with PBS / 0.05% Tween 20, and then blocked with PBS / 2% milk powder. Diluted murine serum samples, collected from each Histopaque separation, were further diluted to 1:1600. These were aliquoted in triplicate (100 μl per well) and incubated for 1 hour at 37°C. Horse radish peroxidase (HRP) conjugated anti-mouse IgG secondary antibody (Abcam, ab99774) was then added (1:5000; 100 μl per well), and the plates incubated for a further 1 hour. After the addition of HRP substrate (80 μl per well) (Stabilised TMB, Life Technologies), the plates were incubated at room temperature in the dark for 5 minutes and read using a FLUOstar Omega plate reader (BMG LABTECH), after the addition of 40 μl 1M HCl.

### Statistics

Analyses were performed in GraphPad PRISM v8.0. S.D. Background cellularity and CD45^+^, CD4^+^ and CD8^+^ cut-offs were set as mean + 3 × S.D. Data sets were compared using a 2-sample t-test with Welch correction. If data were not normally distributed, as assessed using a Shapiro-Wilk test, a Mann-Whitney rank sum test was used.

## Supplementary Figures

**Figure 1 - figure supplement 1.**
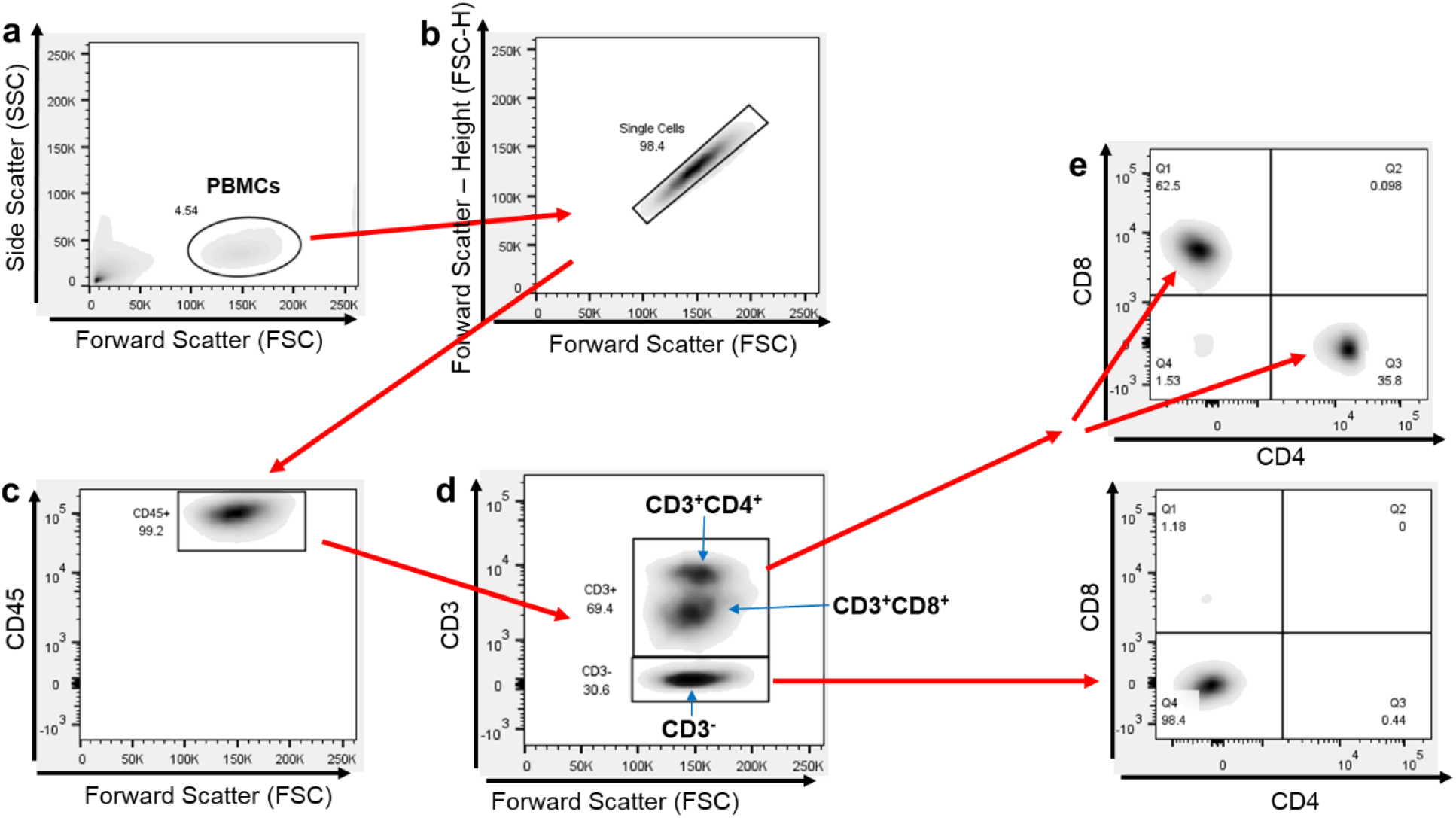
Flow cytometry gating strategy. **a.** PBMCs isolated in the black oval based on forward (FSC) and side (SSC) scatter spectral properties. **b.** Singlets isolated. **c.** Population staining +ve with anti-CD45 antibody. **d.** CD45^+^ population separated by CD3 positivity. **e.** Both CD3^+^ and CD3^-^ populations separated by CD4 and CD8 markers.

**Figure 4 - figure supplement 1.**
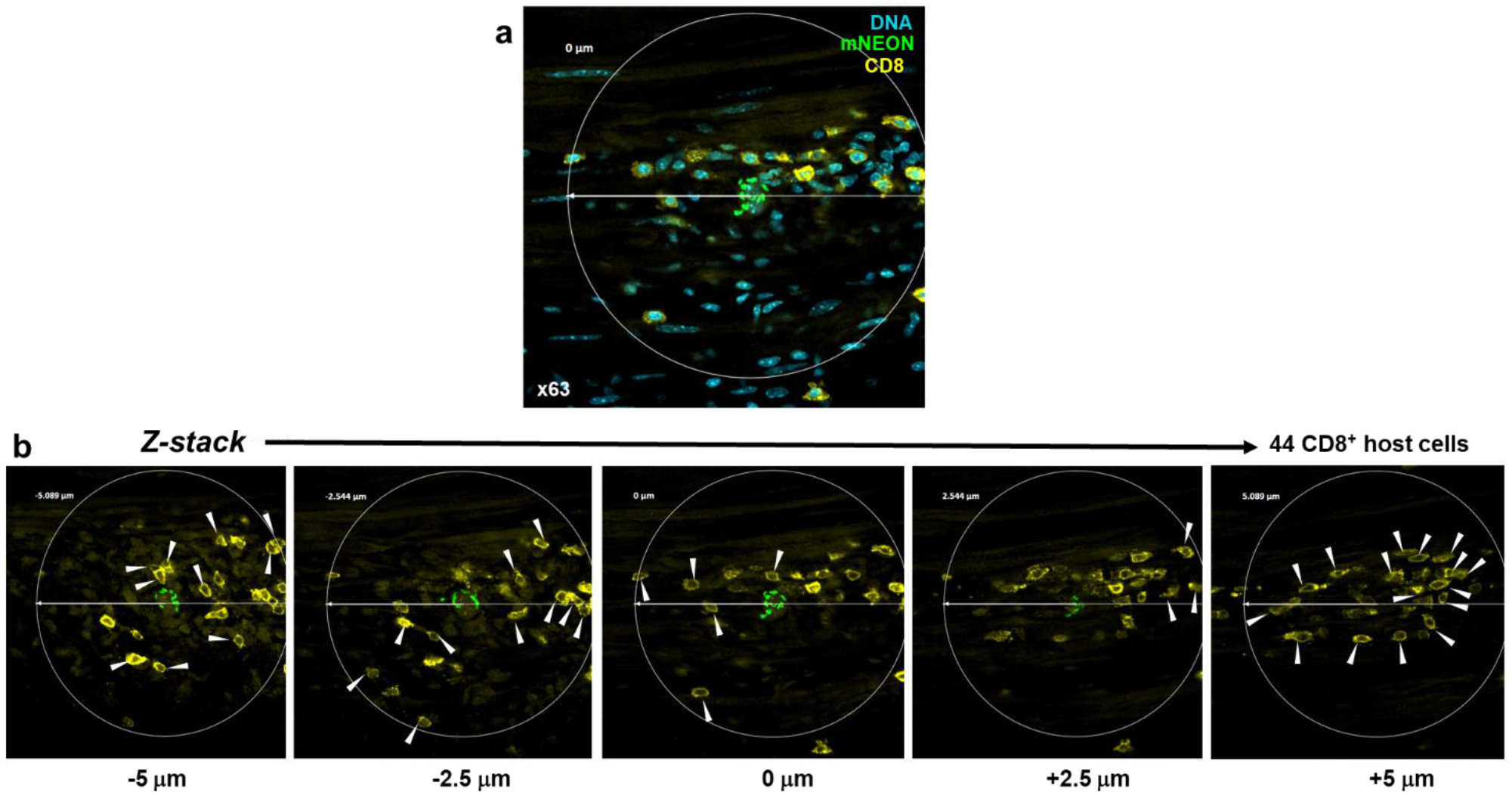
Establishing the extent of CD8^+^ T cell recruitment to infection foci using 3-dimensional serial Z-stack confocal imaging. **a.** A parasite nest detected in the whole mounted colonic gut wall of a mouse chronically infected with *T. cruzi* CL Luc::mNeon (Materials and Methods). Parasites, green; DNA, blue (DAPI); CD8^+^ T cells, yellow (stained with antibody prior to mounting). The area selected for Z-stack imaging is identified by a 200 μm diameter circle, centred on the parasite nest. **b.** The local density of CD8^+^ host cells was determined by counting the number of stained cells (yellow) in a series of Z-stack images acquired with a Zeiss LSM880 confocal microscope from 5 μm above and below the centre of the parasite nest on the Z-axis, a cylinder volume of 314 μm^3^. Any cells that fell within the 200 μm diameter circle were included. The number of hyper-local CD8^+^ T cells was calculated to be 44.

**Figure 4 - figure supplement 2.**
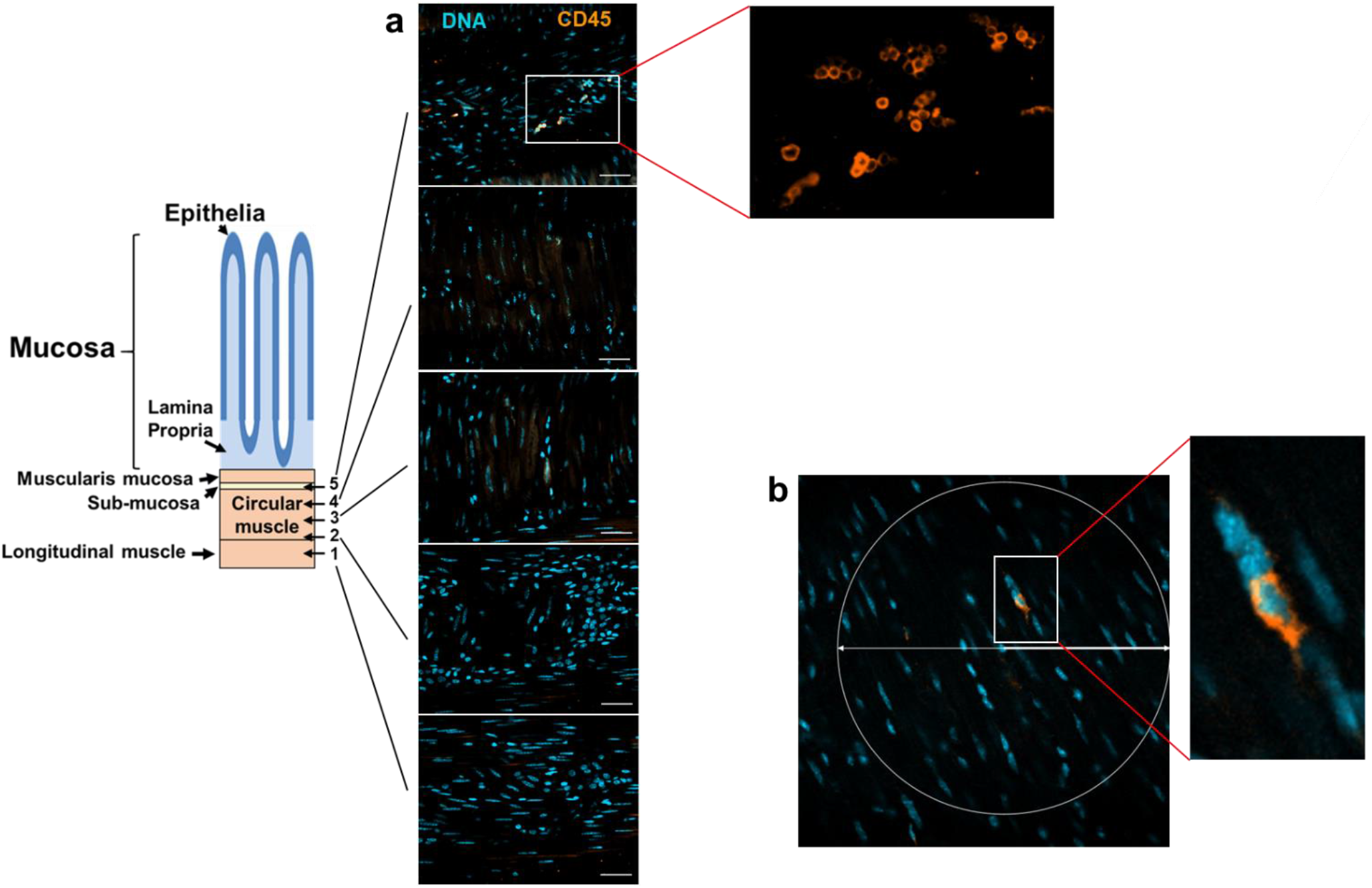
The longitudinal and transverse smooth muscle layers of the colon are largely devoid of CD45^+^ leukocytes in non-infected C3H/HeN mice. **a.** Serial Z-stack images of a whole mounted colonic gut wall from an age-matched non-infected C3H/HeN mouse. DNA, blue (DAPI); CD45^+^, orange. Scale bars=20 μm. The images correspond to the cross-sectional regions of the colon indicated in the schematic (1–5). CD45^+^ cells can be readily detected in the submucosal layer (inset). **b.** Rare example of a CD45^+^ cell within the longitudinal and transverse smooth muscle layers. A 200 μm diameter circle is superimposed on the image.

