## Supplementary data for "Incomplete recruitment of protective T cells facilitates *Trypanosoma cruzi* persistence in the mouse colon"

### Supplementary Figures

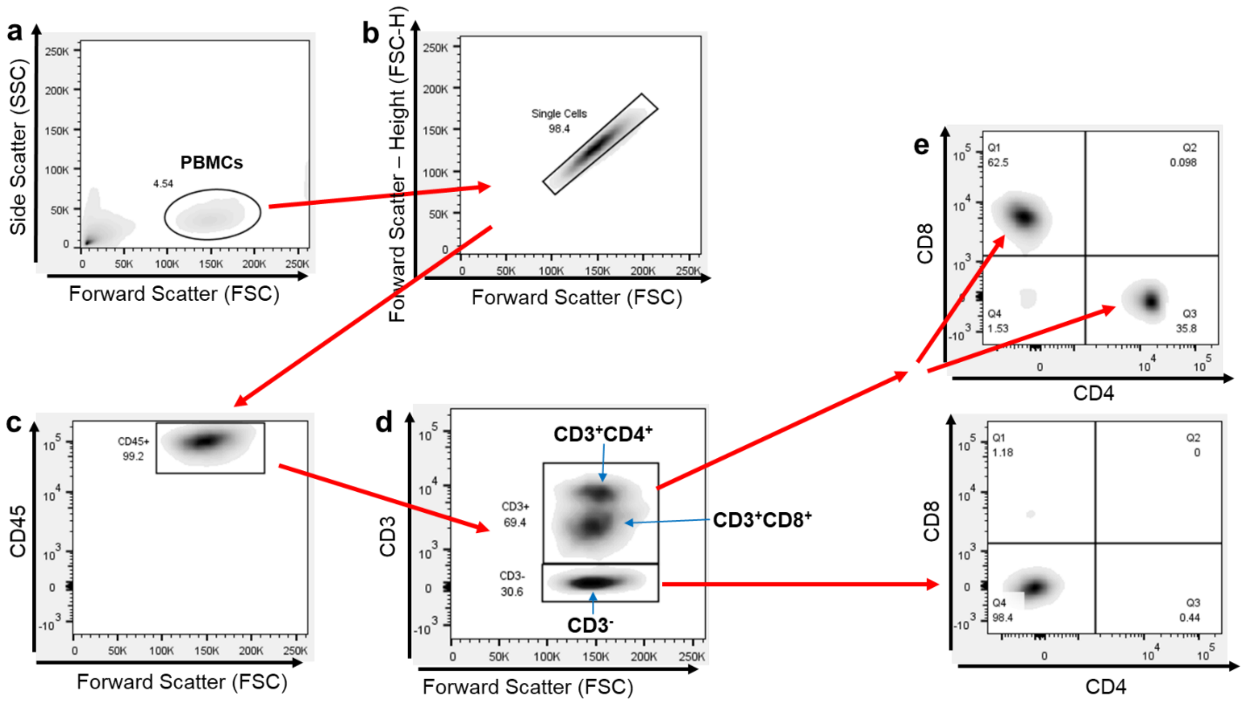


**Figure 1 – figure supplement 1. Flow cytometry gating strategy. a.** PBMCs isolated in the black oval based on forward (FSC) and side (SSC) scatter spectral properties. **b.** Singlets isolated. **c.** Population staining +ve with anti-CD45 antibody. **d.** $CD45^+$ population separated by CD3 positivity. **e.** Both $CD3^+$ and $CD3^-$ populations separated by CD4 and CD8 markers.

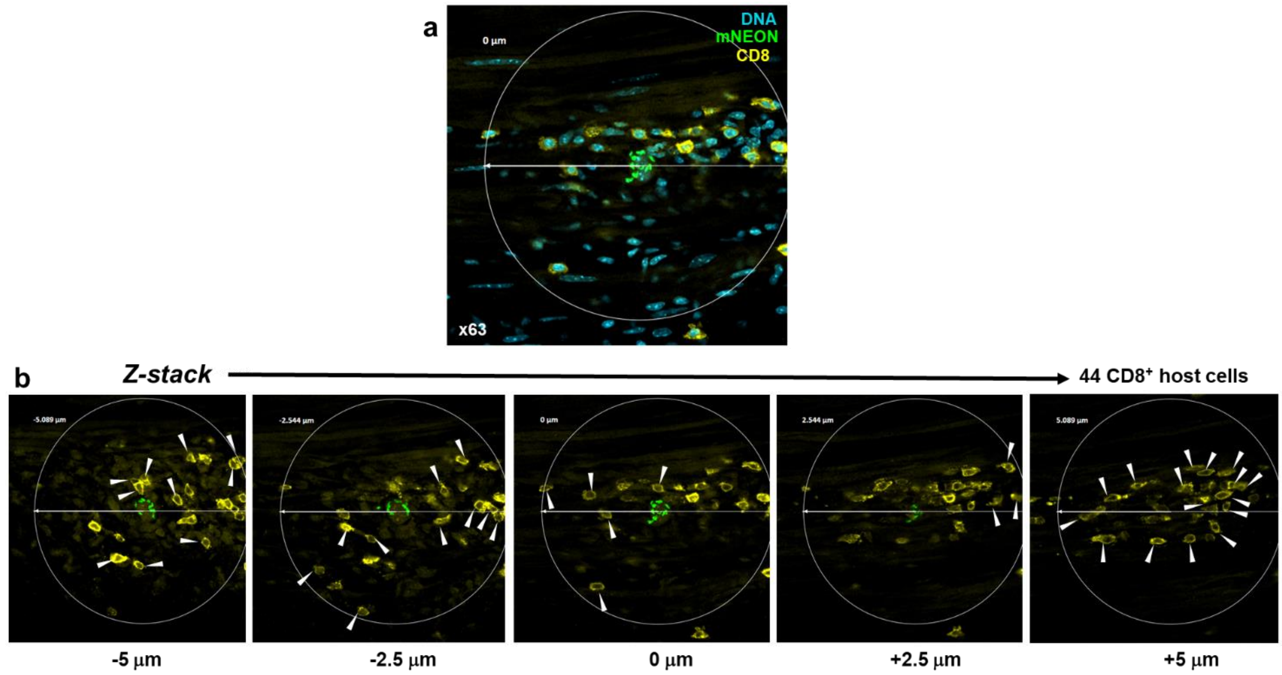


**Figure 4 - supplement figure 1. Establishing the extent of CD8$^+$ T cell recruitment to infection foci using 3-dimensional serial Z-stack confocal imaging. a.** A parasite nest detected in the whole mounted colonic gut wall of a mouse chronically infected with *T. cruzi* CL Luc::mNeon (Materials and Methods). Parasites, green; DNA, blue (DAPI); CD8$^+$ T cells, yellow (stained with antibody prior to mounting). The area selected for Z-stack imaging is identified by a 200 µm diameter circle, centred on the parasite nest. **b.** The local density of CD8$^+$ host cells was determined by counting the number of stained cells (yellow) in a series of Z-stack images acquired with a Zeiss LSM880 confocal microscope from 5 µm above and below the centre of the parasite nest on the Z-axis, a cylinder volume of 314 µm$^3$. Any cells that fell within the 200 µm diameter circle were included. The number of hyper-local CD8$^+$ T cells was calculated to be 44.

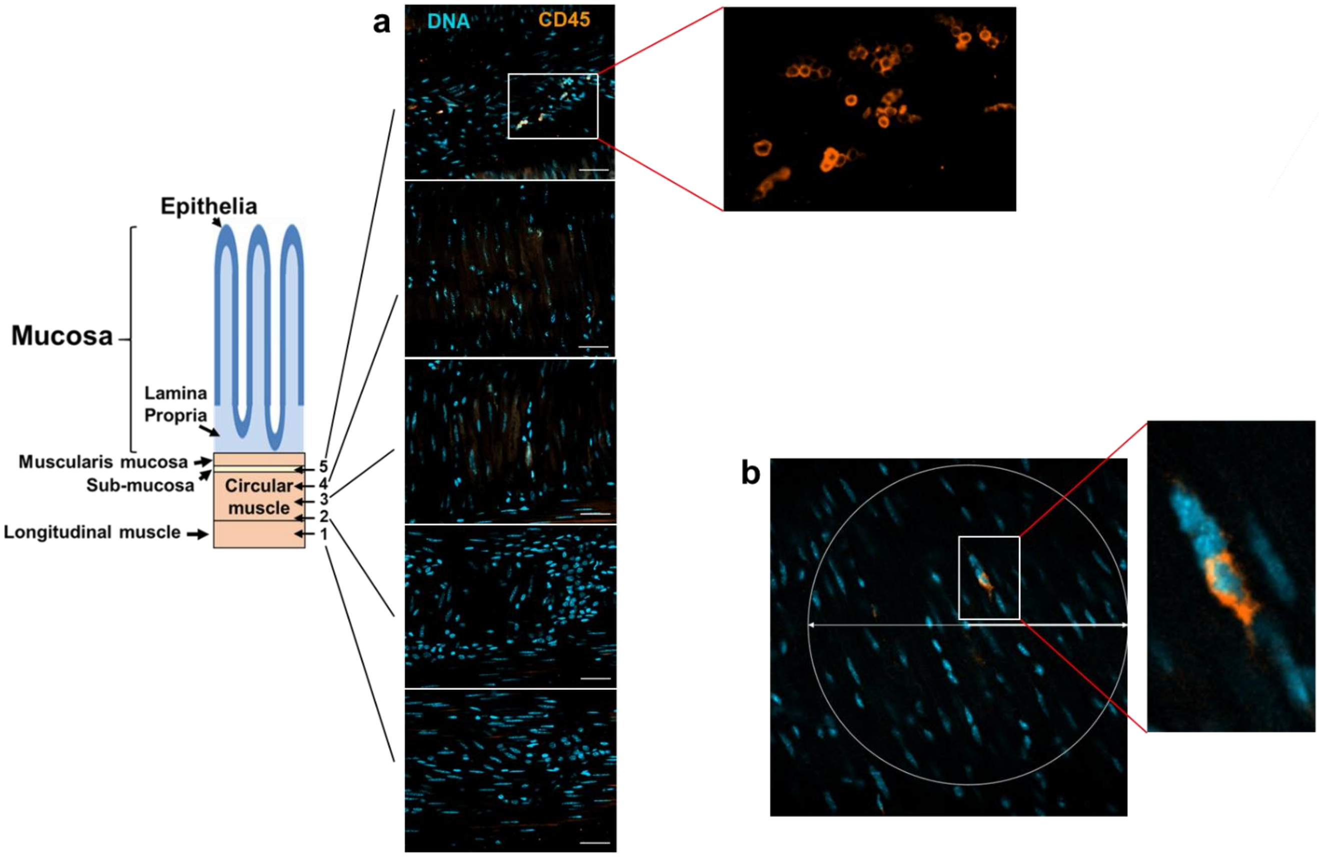


**Figure 4 - supplement figure 2. The longitudinal and transverse smooth muscle layers of the colon are largely devoid of CD45⁺ leukocytes in non-infected C3H/HeN mice. a.** Serial Z-stack images of a whole mounted colonic gut wall from an age-matched non-infected C3H/HeN mouse. DNA, blue (DAPI); CD45⁺, orange. Scale bars=20 μm. The images correspond to the cross-sectional regions of the colon indicated in the schematic (1-5). CD45⁺ cells can be readily detected in the sub-mucosal layer (inset). **b.** Rare example of a CD45⁺ cell within the longitudinal and transverse smooth muscle layers. A 200 μm diameter circle is superimposed on the image.
